# SARS-CoV-2 spike D614G variant confers enhanced replication and transmissibility

**DOI:** 10.1101/2020.10.27.357558

**Authors:** Bin Zhou, Tran Thi Nhu Thao, Donata Hoffmann, Adriano Taddeo, Nadine Ebert, Fabien Labroussaa, Anne Pohlmann, Jacqueline King, Jasmine Portmann, Nico Joel Halwe, Lorenz Ulrich, Bettina Salome Trüeb, Jenna N. Kelly, Xiaoyu Fan, Bernd Hoffmann, Silvio Steiner, Li Wang, Lisa Thomann, Xudong Lin, Hanspeter Stalder, Berta Pozzi, Simone de Brot, Nannan Jiang, Dan Cui, Jaber Hossain, Malania Wilson, Matthew Keller, Thomas J. Stark, John R. Barnes, Ronald Dijkman, Joerg Jores, Charaf Benarafa, David E. Wentworth, Volker Thiel, Martin Beer

## Abstract

During the evolution of SARS-CoV-2 in humans a D614G substitution in the spike (S) protein emerged and became the predominant circulating variant (S-614G) of the COVID-19 pandemic^1^. However, whether the increasing prevalence of the S-614G variant represents a fitness advantage that improves replication and/or transmission in humans or is merely due to founder effects remains elusive. Here, we generated isogenic SARS-CoV-2 variants and demonstrate that the S-614G variant has (i) enhanced binding to human ACE2, (ii) increased replication in primary human bronchial and nasal airway epithelial cultures as well as in a novel human ACE2 knock-in mouse model, and (iii) markedly increased replication and transmissibility in hamster and ferret models of SARS-CoV-2 infection. Collectively, our data show that while the S-614G substitution results in subtle increases in binding and replication *in vitro*, it provides a real competitive advantage *in vivo*, particularly during the transmission bottle neck, providing an explanation for the global predominance of S-614G variant among the SARS-CoV-2 viruses currently circulating.

## Main Text

In late 2019, severe acute respiratory syndrome coronavirus 2 (SARS-CoV-2) emerged in Wuhan, Hubei province, China^2,3^ and rapidly developed into the COVID-19 pandemic. By the end of September 2020, the worldwide death toll had passed one million people with more than 37 million infections^4^. Symptoms are usually mild; however in more vulnerable groups, such as aged individuals or people with comorbidities, SARS-CoV-2 can cause life-threatening pneumonia^5^. Cell entry of SARS-CoV-2 is dependent on the interaction of the spike glycoprotein (S) and the host cell surface receptor angiotensin-converting enzyme 2 (ACE2)^3,6^. S is a homotrimeric class I fusion protein consisting of two subunits S1 and S2, which are separated by a protease cleavage site. The S1 forms a globular head and is essential for receptor binding, while S2 is responsible for fusion of the viral envelope with host cell membranes. During the entry process, the receptor-binding domain (RBD) within the S1 subunit binds ACE2, generating conformational changes in the S2 subunit, which facilitates virus internalization^7,8^. S-D614G is a protein variant containing a substitution in the S protein outside of the RBD and is thought to cause a conformational change. It is believed to weaken the interprotomer latch in the S protein trimer between the S1 and S2 domains and causes a more “open” conformation that improves ACE2 binding and increases the probability of infection ^1,9^. Over the course of the pandemic, the SARS-CoV-2 S-614G variant rapidly superseded the parental S-614D variant in frequency to become globally dominant. Such a shift in genotype frequency might be caused by a founder effect following introduction into a highly interconnected population. However, there are multiple lines of evidence suggesting that the S-614G variant may confer a fitness advantage compared to S-614D: increased frequency of S-614G in distinct geographical regions, initial experimental evidence with pseudotyped lentiviruses^9^ or vesicular stomatitis viruses ^8^, and reports of the S-614G variant being associated with higher viral loads^1^. To better address the role that the S-D614G substitution has played in the dissemination and predominance of this SARS-CoV-2 variant during the COVID-19 pandemic, we characterized S protein binding to human ACE2 (hACE2) and replication kinetics *in vitro*, and evaluated infection and transmission dynamics *in vivo* using three different animal models. The data show that the S-D614G substitution confers increased binding to the hACE2 receptor and increased replication in primary human airway epithelial cultures. Moreover, comparison of recombinant isogenic SARS-CoV-2 variants demonstrates that S-614G substitution provides competitive advantage in a hACE2 knock-in mouse model, and markedly increases replication and transmission in Syrian hamster and ferret models.

## Results

### SARS-CoV-2 S-614G binds to hACE2 more efficiently

To determine whether the S-D614G substitution directly affects the binding between the S and hACE2, we first used the biolayer interferometry (BLI) technology to quantify their biding affinity. Because the S1 component of the S is the domain that interacts with receptor, a reductionist approach was used to determine if the D614G played a role in hACE2 binding by monomeric S1 proteins. Pre-biotinylated polyhistidine-tagged S1 proteins with 614D or 614G (S1-614D and S1-614G, respectively) both bind efficiently to hACE2; however, S1-614G (KD = 1.65 nM) showed about 2-fold higher affinity than S1-614D (KD = 3.74 nM) (Figure 1A). When the full-length monomeric spike ectodomain was used in the assay, the S-614G protein also showed higher affinity to hACE2 than S-614D (Extended Data Figure 1A). The enhanced binding to hACE2 protein rendered by the S-D614G substitution also resulted in enhanced S1 binding to Baby Hamster Kidney (BHK) cells expressing exogenous hACE2 (BHK-hACE2) in a different binding assay (Figure 1B, Extended Data Figure 1B). We incubated polyhistidine-tagged S1-614D or S1-614G proteins with BHK-hACE2 and analyzed the binding efficiency of S1 to the cells using flow cytometry. At the same S1 concentration, more S1-614G bound to the BHK-hACE2 cells than S1-614D (Figure 1B, Extended Data Figure 1B). Recombinant S1 constructs that express two S1 molecules attached to an IgG carboxyl-terminus were generated to further evaluate the impact the S-D614G substitution. An even more striking difference was observed with the pair of Fc(IgG)-tagged S1-614D or S1-614G proteins used for binding studies instead of polyhistidine-tagged S1 constructs (Figure 1B, Extended Data Figure 1B).

**Figure 1.**
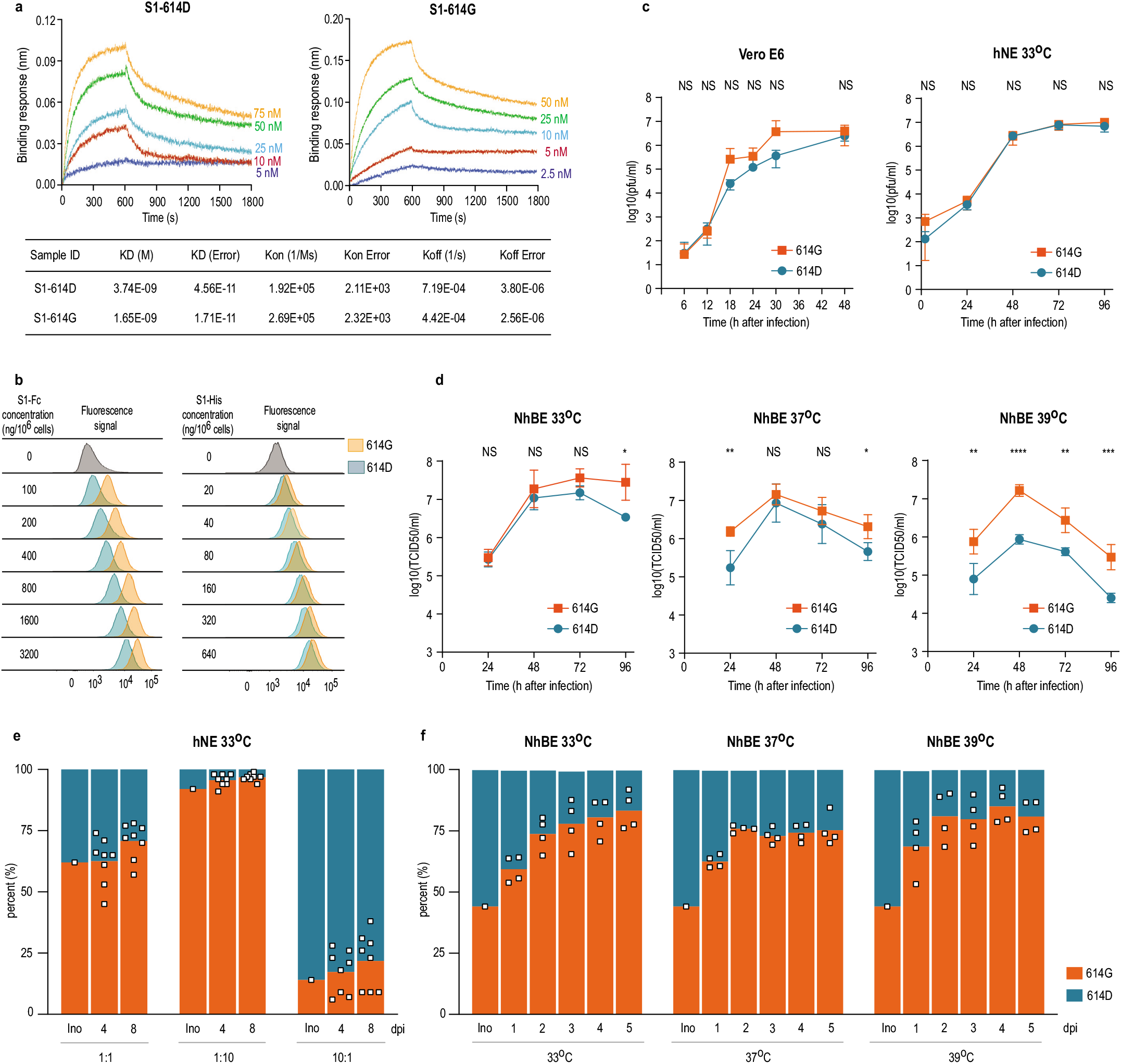
In vitro characterization of S1 proteins and recombinant SARS-CoV-2^S-614D^ and SARS-CoV-2^S-614G^ viruses. **(a)** Affinity between S1 and hACE2 determined by Bio-layer interferometry. Biotinylated S1 protein (S1-614D or S1-614G) was loaded onto surface of streptavidin biosensors. Association was conducted using hACE2 protein followed by dissociation. (**b)** Binding of Fc-tagged or polyhistidine-tagged S1 to BHK-hACE2 cells is shown as peaks of fluorescence detected by flow cytometry. (**c)** Replication kinetics of recombinant viruses in (left) Vero E6 at 37°C and (right) hNE at 33°C. Supernatant was collected at indicated time points and titrated by plaque assay. Data represent the mean ± s.d. of three replicates (Vero E6) and four replicates (hNE). (**d)** Replication kinetics of recombinant viruses in NhBE at 33°C (left), 37°C (middle) and 39°C (right). NhBE were infected with 100,000 PFU of each virus. Supernatants were collected daily and titrated by TCID50 assay. Data represent the mean ± s.d. of four replicates. (**c-d**) Statistical significance was determined by two-sided unpaired Student’s *t*-test without adjustments for multiple comparisons. (**c**) *P* values (left to right): left, NS, *P*=0.9132; NS *P*=0.0604; NS *P*=0.2394; NS *P*=0.2389; NS *P*=0.2778; NS *P*=0.2781; right, NS *P*=0.1520; NS *P*=0.3891; NS *P*=0.9110; NS *P*=0.8985; NS *P*=0.1464. (**d**) *P* values (left to right): left, NS *P*=0.7943; NS *P*=0.5025; NS *P=*0.6683; NS *P*=0.8985; \**P*=0.0220; middle, \*\**P*=0.0065; NS *P*=0.4660; NS *P*=0.3134; \**P*=0.0159; right, \*\**P*=0.0094; \*\*\*\**P*<10^−4^; \*\**P*=0.0028; \*\*\**P*=0.0009. (**e-f)** Competition assay of recombinant viruses in hNE at 33°C and NhBE at 33°C, 37°C and 39°C. The inoculum was prepared by mixing two viruses at 1:1 ratio based on PFU ml^−1^ and used for infection of hNE and NhBE. Apical wash and supernatant were collected daily, and extracted RNA was used for sequencing. (**e-f**) Bar graph shows proportion of sequencing reads encoding either S-614D or S-614G, and square dots represent individual data points.

### Increased replication of SARS-CoV-2^S-614G^ virus in primary human epithelial cells

To assess the impact of S-614G in the context of virus infection we generated an isogenic D614G virus pair based on our reverse genetics system for SARS-CoV-2^10^. The molecular clone is based on the Wuhan-Hu-1 isolate possessing the S-614D variant (SARS-CoV-2^S-614D^) ^10,11^. The sequence of the isogenic S-614G variant was engineered to have an A to G nucleotide change at position 23,403 to encode a glycine at the S protein position 614. The identity of the resulting recombinant SARS-CoV-2^S-614G^ variant was confirmed by full-length sequencing from the passage 1 virus stock that was used for subsequent experiments. Replication kinetics of SARS-CoV-2^S-614D^ and SARS-CoV-2^S-614G^ in Vero E6 cells marginally differed (Figure 1C). We assessed replication kinetics in primary human nasal epithelial (hNE) and primary normal human bronchial epithelial (NhBE) cultures that were grown under air-liquid interface conditions and resemble the pseudostratified epithelial lining of the human respiratory epithelium. No significant difference in the primary hNE cells following infection of SARS-CoV-2^S-614D^ or SARS-CoV-2^S-614G^ at 33°C, the temperature of the nasal epithelium was observed (Figure 1C). In contrast, SARS-CoV-2^S-614G^ displayed elevated titers in primary NhBE cells at temperatures of 33°C, 37°C and 39°C, that resemble temperatures of the upper and lower respiratory tract, or fever, respectively (Figure 1D). Similarly, infection kinetics of NhBE cells with natural isolates SARS-CoV-2/USA-WA1/2020 (USA-WA1, S-614D) or SARS-CoV-2/Massachusetts/VPT1/2020 (MA/VPT1, S-614G) revealed increased titers for the S-614G variant (Extended Data Figure 1C). To refine this analysis, we performed competition experiments by infecting hNE and NhBE cultures with a mixture of both viruses, SARS-CoV-2^S-614D^ and SARS-CoV-2^S-614G^, at defined ratios. In both primary human respiratory culture systems, the ratio of 614G:614D shifted in favor of SARS-CoV-2^S-614G^ during five or eight days of infection (Figure 1E, 1F, Extended Data Figure 1D). Collectively, these results show that the D614G change in the S protein is associated with enhanced hACE2 binding and increased replication in primary human airway epithelial models of SARS-CoV-2 infection.

### Increased replication of SARS-CoV-2^S-614G^ in hACE2 knock-in mice

Mice do not support efficient replication of SARS-CoV-2 unless they are genetically engineered to express hACE2^12,13^. To evaluate the relative fitness of the SARS-CoV-2^S-614G^ variant *in vivo*, we generated knock-in mice expressing the authentic SARS-CoV-2 human receptor *hACE2* under the endogenous regulatory elements of the mouse *Ace2* gene (hACE2-KI, Extended Data Figure 2A). Eight heterozygous female mice were inoculated intranasally (i.n.) in a competition experiment with a mixture of both viruses, SARS-CoV-2^S-614D^ and SARS-CoV-2^S-614G^, using 1×10^5^ plaque forming unit (PFU) of each variant (Figure 2A). Viral RNA loads were monitored daily in oropharyngeal swabs and in various organs and tissues by real-time PCR at days 2 and 4 post infection (p.i.). No significant body weight loss in hACE2-KI mice up to day 4 p.i. were observed (Extended Data Figure 2B). Longitudinal analysis of daily oropharyngeal swabs revealed efficient virus replication in the upper respiratory tract of hACE2-KI mice (Figure 2B). Accordingly, tissue samples collected at day 2 and 4 p.i. revealed high viral RNA loads in the nasal conchae, lungs and olfactory bulbs and at lower levels in the brain (Extended Data Figure 2C). Low to undetectable levels of virus were observed in spleen, small intestine, kidneys and feces (data not shown). No overt pathological lesions were found in the lungs of hACE2-KI compared to wild-type mice at day 2 and 4 p.i. (Extended Data Table 1, 2). Sequencing analysis of the oropharyngeal swabs revealed a net advantage for the SARS-CoV-2^S-614G^ variant over SARS-CoV-2^S-614D^ variant in most animals and time points (Figure 2C). In the organs, a similar replication advantage was found for the SARS-CoV-2^S-614G^ variant (Figure 2D). Collectively, these results demonstrate increased replication of SARS-CoV-2^S-614G^ in a mouse model of SARS-CoV-2 infection in the context of the expression of the authentic human receptor *hACE2*.

**Figure 2.**
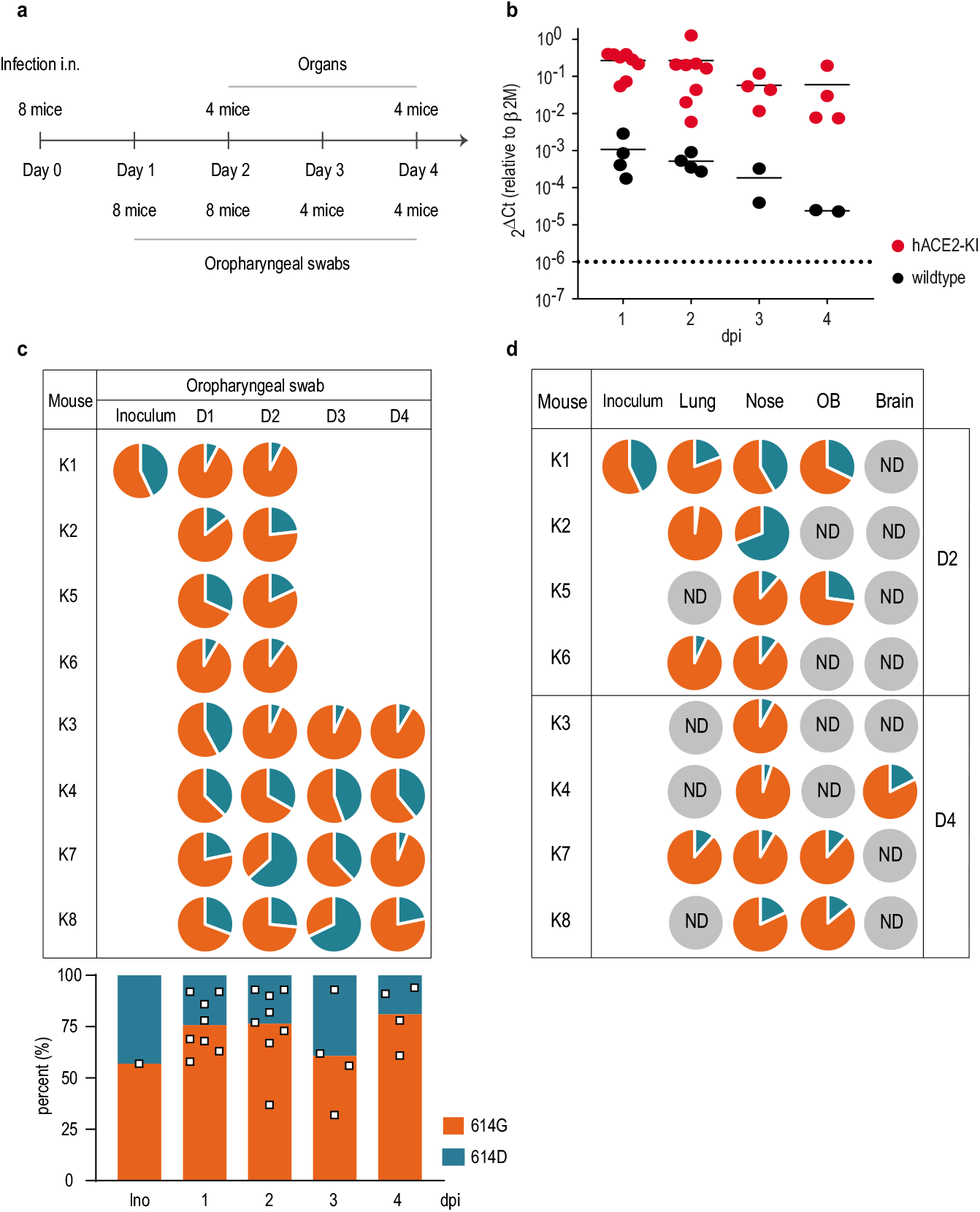
Replication of SARS-CoV-2^S-614D^ and SARS-CoV-2^S-614G^ viruses in hACE2 knock-in mice. (**a**) Experimental scheme for infection of hACE2-KI mice intranasally infected recombinant SARS-CoV-2^S-614D^ and SARS-CoV-2^S-614G^ viruses. Oropharyngeal swabs were sampled daily and tissue samples were analyzed in sub-groups of 4 mice at 2 and 4 days post infection (dpi) in two independent experiments. (**b**) Quantitative RT-PCR analysis of oropharyngeal swabs of inoculated hACE2-KI and wild-type mice. (**c,d**) Pie chart representation of mean frequencies of A or G nucleotide at position 23,403 corresponding to SARS-CoV-2^S-614D^ and SARS-CoV-2^S-614G^, respectively. Each pie chart illustrates the ratio of A/G detected from individual oropharyngeal swab samples (c) and tissues (d) at indicated time post infection. OB, olfactory bulb; ND, not detected.

### SARS-CoV-2^S-614G^ displays increased replication and transmissibility in hamsters and ferrets

Hamsters are highly susceptible to SARS-CoV-2 infection and develop disease that closely resembles pan-respiratory, fulminant COVID-19 disease in humans^14,15^. In contrast, in ferrets SARS-CoV-2 primarily replicates within the upper respiratory tract, resembling mild human infections. However, both animal models efficiently reflect transmission events by direct contact. By using a competition experimental approach *in vivo*, as shown for the hACE2-KI mice, numerical dominance of one recombinant variant should be the result of relevant advantages.

Therefore, direct “one-to-one” transmission experiments were conducted. Six donor Syrian hamsters were inoculated i.n. at equal ratios with SARS-CoV-2^S-614D^ and SARS-CoV-2^S-614G^ using 1×10^4.77^ TCID_50_/animal (calculated from back titration of the original material). Analysis of the mixed inoculum by amplicon sequencing and absolute quantification using allele specific locked nucleic acid (LNA) probes confirmed similar viral RNA ratios of both variants (Figure 3). At 24 hours after inoculation, each donor was cohoused with one naive hamster. Weight changes, as well as clinical signs were monitored and nasal washes were collected daily. Viral RNA load in nasal washings, and changes in body weight, were congruent to previously published data^14,15^ (Extended Data Figure 3A, B). Analysis of the viral RNA sequence composition of nasal washings revealed a prominent shift towards SARS-CoV-2S-614G within 48 hours post inoculation (Figure 3). Transmission efficiency was one hundred percent, and analysis of the RNA sequence composition showed that the SARS-CoV-2^S-614G^ variant represented >90% of the viral RNA in the contact animals (Figure 3). In summary, hamsters inoculated with both variants, SARS-CoV-2^S-614D^ and SARS-CoV-2^S-614G^, in an equal ratio, transmit primarily SARS-CoV-2^S-614G^.

**Figure 3.**
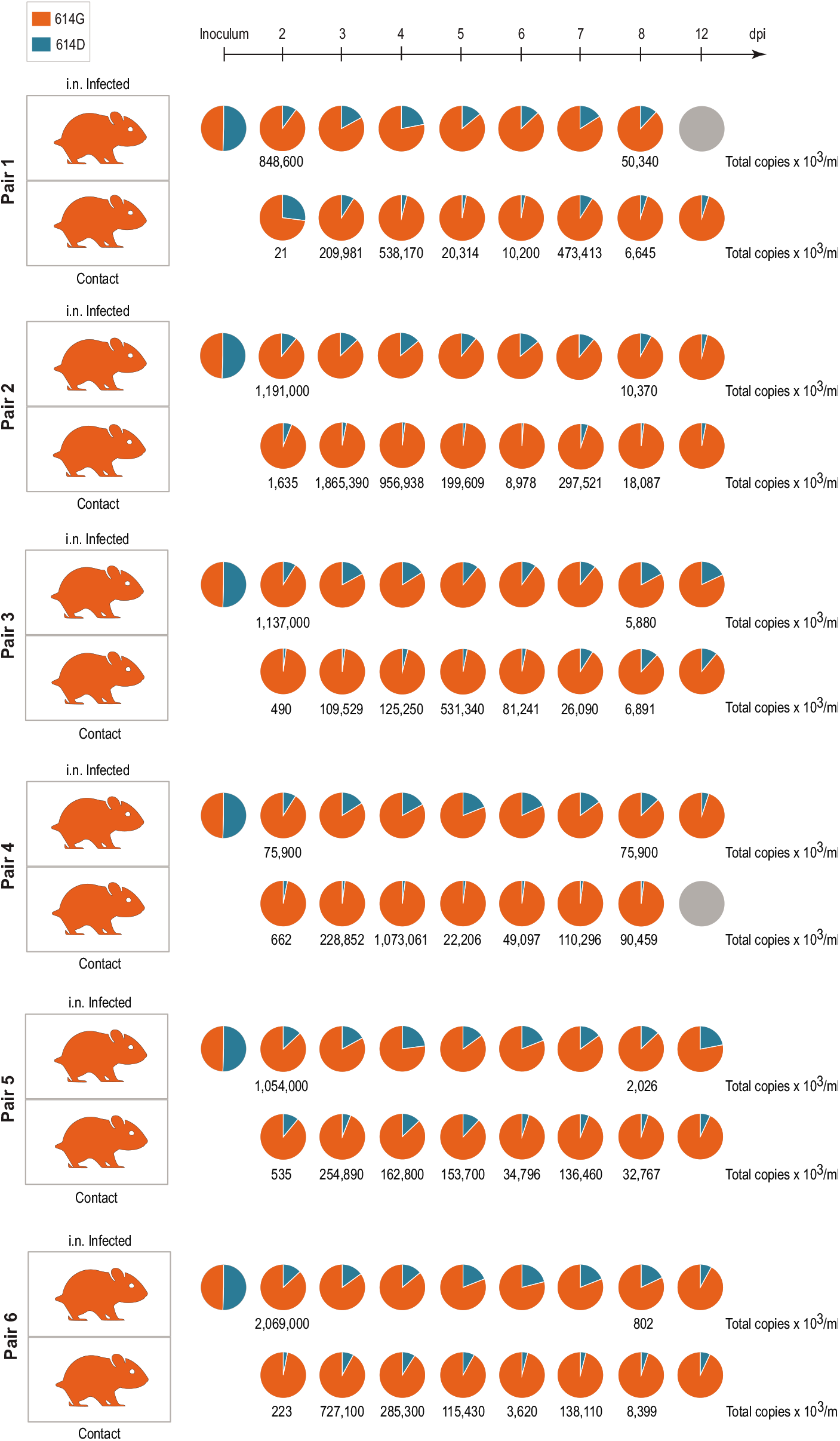
Replication and transmission of SARS-CoV-2^S-614D^ and SARS-CoV-2^S-614G^ viruses in Syrian hamsters. Transmission of SARS-CoV-2^S-614D^ and ^S-614G^ variant by hamsters in a pairwise one-by-one setup with direct contact of donor and cohoused contact hamsters is illustrated. Samples of nasal washings were taken daily between days 2 to 8 post infection (dpi) and finally at 12 dpi and were analyzed. Pie chart representation of fraction of A or G nucleotide at position 23,403 corresponding to SARS-CoV-2^S-614D^ and SARS-CoV-2^S-614G^, respectively, measured by amplicon sequencing. Genome copies were calculated from RT-qPCR using a standard RNA. Orange coloring of the hamster silhouette refer to detection of G (SARS-CoV-2^S-614G^), while blue coloring indicates detection of A (SARS-CoV-2^S-614D^) on most time points. Grey coloring signals no infection detected

To exclude possible differences in hamster’s affinity to one variant or divergence of kinetics of the replication cycle six hamsters were inoculated i.n. with either SARS-CoV-2^S-614D^ (1× 10^5.1^ TCID_50_/animal, calculated from back titration of the original material), or SARS-CoV-2^S-614G^ (10^4.5^ TCID_50_/animal, calculated from back titration of the original material) and were monitored for four consecutive days. No marked differences in body weights, titers of shed virus, or viral loads in respiratory tract tissue were observed between the two groups in the acute phase (Extended Data Figure 3C-E). For both variants, highest genome loads were found in the nasal conchae, followed by pulmonary tissue (Extended Data Figure 3E). These observations confirm that in the case of SARS-CoV-2^S-614D^ or SARS-CoV-2^S-614G^ infections, the S-D614G substitution does not seem crucial for clinical outcomes, which again underscores the hamster as a highly sensitive disease model. Rather, the advantage of the SARS-CoV-2^S-614G^ variant over the SARS-CoV-2^S-614D^ has to be adequately large to fully suppress the latter variant within a single *in vivo* replication cycle, which accurately reflects the evolution of SARS-CoV-2^S-614G^ in humans during the COVID-19 pandemic.

Since ferrets are a good transmission model ^18^, we performed a direct “one-to-one” transmission experiment using an equal mixture of the isogenic SARS-CoV-2 variants. Six animals were intranasally inoculated with the mix of SARS-CoV-2^S-614D^ and SARS-CoV-2^S-614G^ (10^5.4^ TCID_50_/animal calculated from back titration of the original material). For all six inoculated ferrets SARS-CoV-2-infection could be confirmed, and both body weight changes and viral RNA loads in nasal washings (Figure 4, Extended Data Figure 4A, B) reflected published data^16,17^. In five of the six inoculated ferrets, SARS-CoV-2^S-614G^ became the dominant variant (Figure 4). In addition, SARS-CoV-2 transmission occurred in four of the six ferret pairs, and in each pair with successful transmission the S-614G variant prevailed over S-614D (Figure 4). All amplicon sequencing data of the ferret samples were also retested by absolute quantification using allele specific locked nucleic acid (LNA) probes and digital PCR analysis.

**Figure 4.**
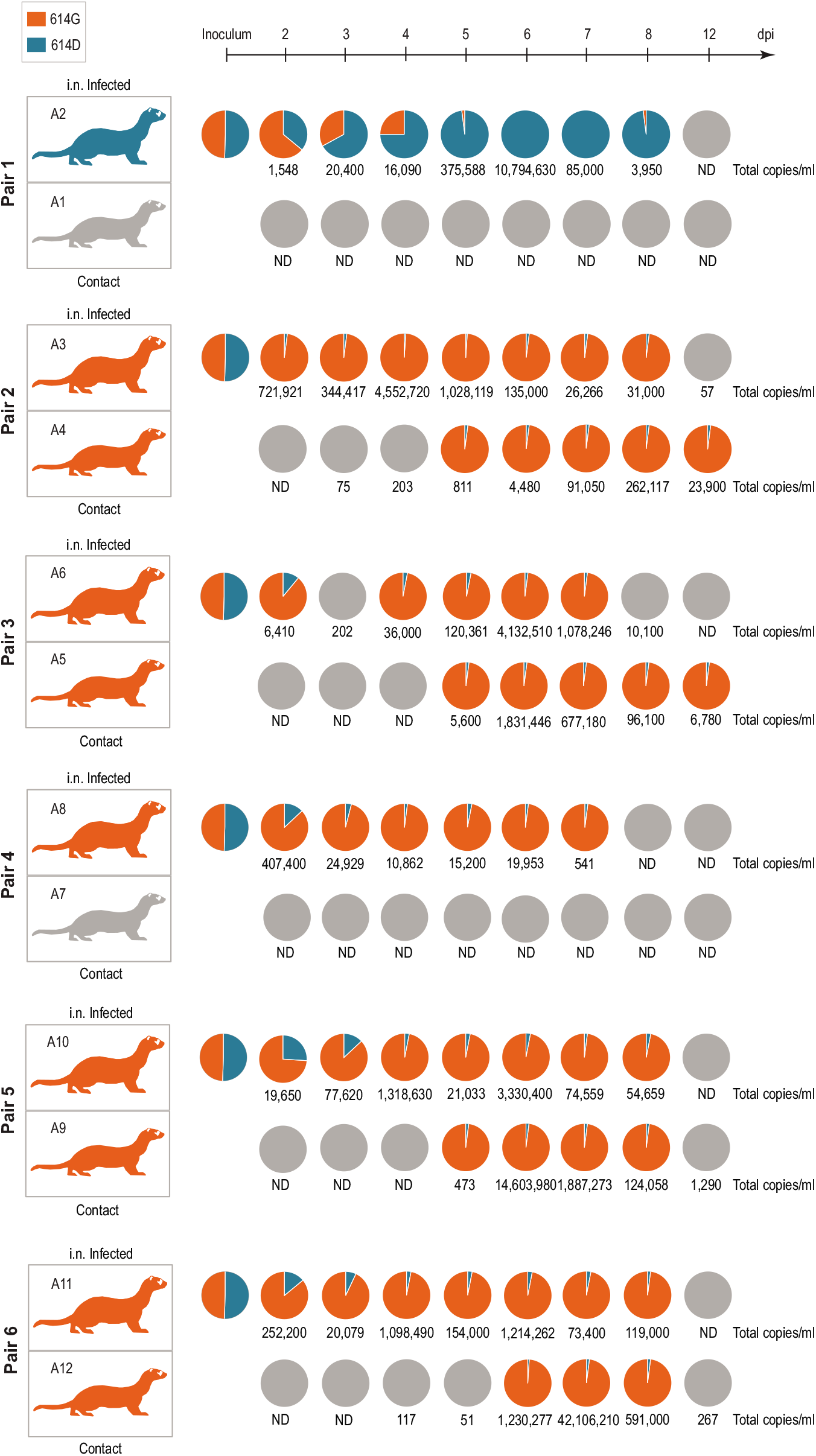
Replication and transmission of SARS-CoV-2^S-614D^ and SARS-CoV-2^S-614G^ viruses in ferrets. Schematic illustration of the experimental setup with six pairs of donor ferrets cohoused with naïve contact ferrets. Samples of nasal washings were taken daily between days 2 to 8 post infection (dpi) and finally at 12 dpi and were analyzed. Pie chart representation of fraction of A or G nucleotide at position 23,403 corresponding to SARS-CoV-2^S-614D^ and SARS-CoV-2^S-614G^, respectively. Each pie chart illustrates the ratio of A/G detected from individual nasal washing samples over time. Orange coloring of the ferret silhouette refer to detection of G (SARS-CoV-2^S-614G^) on most time points, while blue coloring indicates detection of A (SARS-CoV-2^S-614D^). Numbers represent total genome copies ml^−1^ and grey coloring signals no infection or viral genome number too low for A/G ratio determination in sequencing. ND, not detected.

Notably, the donor of pair one with the dominance of SARS-CoV-2^S-614D^ did not transmit despite a high viral genome peak load of more than 10 million copies per ml. In contrast, the lack of transmission in pair 4 where SARS-CoV-2^S-614G^ became the dominant variant is connected to peak viral loads of below 500,000 genome copies per ml (Figure 4). In summary, the competition experiment in ferrets revealed that the SARS-CoV-2^S-614G^ variant preferentially infected and replicated in five out of six inoculated animals, and transmission events succeeded exclusively with the SARS-CoV-2^S-614G^ variant.

## Discussion

SARS-CoV-2 evolution in humans has been proposed to be a non-deterministic process and virus diversification results mainly from random genetic drift, suggesting that there is no strong selective pressure on SARS-CoV-2 in its adaptation to humans^20^. However, since the introduction of the S-D614G change in early 2020, the SARS-CoV-2 S-614G variant has become globally prevalent^1^. A founder effect and a structural change of the SARS-CoV-2 spike protein have been proposed as driving forces establishing the S-614G prevalence. Previous structural studies and the use of pseudotyped viruses have put forward the idea that the S-614G variant may confer increased infectivity, which could be a result of increased “open” RBD conformation for receptor binding as suggested by one study or increased S stability as suggested by another study^1,9^. In contrast to those studies that used recombinant trimeric S, we used a reductionist approach to eliminate the complications due to the “open” or “closed” RBD conformations in trimeric S. We found the S1 or the monomeric S ectodomain with D614G substitution had increased affinity to hACE2, which may be another mechanism underlying the increased replication and transmission of the SARS-CoV-2 D614G variant.

Studies that employed isogenic SARS-CoV-2 D614G variants to assess the phenotype in the context of a SARS-CoV-2 infection were only very recently reported in preprints ^19,20^. Both studies conclude that the SARS-CoV-2 S-614G variant shows increased replication *in vitro* and one study observed earlier transmission in a hamster model. We extended these studies by exploiting various *in vitro* and *in vivo* infection models of SARS-CoV-2, including primary NhBE and hNE cultures, a novel hACE2 knock-in mouse model, a hamster model, and a ferret model. Importantly, throughout these experimental systems we consistently observed an increased fitness of SARS-CoV-2^S-614G^ over SARS-CoV-2^S-614D^ by applying amplicon sequencing techniques as well as allele specific absolute quantification for confirmation. The advantage provided by the D614G change becomes most prominent in competition and transmission experiments in hamsters and ferrets and must therefore be considered as a driving force leading to the global dominance of the SARS-CoV-2 614G variant.

Our data are also in agreement with reported functional changes conferred by the D614G substitution in the S protein^1^ and infections studies using pseudotyped viruses demonstrating increased infection^9,21^. Although our studies establish a phenotype of increased replication and transmission of the SARS-CoV-2 S-614G variant, there is no evidence for a phenotypic change in pathogenicity in an animal model. This is important to state, because infection with the SARS-CoV-2 S-614G variant is not associated with the development of severe COVID-19 in humans^1^.

The ongoing pandemic will likely give rise to additional SARS-CoV-2 variants that may display phenotypic changes and further adaptations to humans. The ability to rapidly trace the genetic variability of emerging variants using whole-genome sequencing, reconstructing emerging virus variants, and assessing their phenotypes will allow rapid response to their emergence with appropriate countermeasures. The development, improvement and characterization of suitable animal models that recapitulate SARS-CoV-2 replication, transmission and pathogenicity in humans will provide a platform to assess the potential implications of these emerging variants. The novel mouse model based on hACE2 expression under the endogenous regulatory elements of the mouse *Ace2* gene will be a valuable tool and will complement existing animal models of SARS-CoV-2 infection. Similarly, as we have shown here, to demonstrate increased replication and transmissibility of SARS-CoV-2^S-614G^, the phenotypic assessment of future pandemic variants will likely require several complementing animal models that together reflect aspects of SARS-CoV-2 replication, transmission and pathogenicity in humans.

## Acknowledgements

This work was supported by the Swiss National Science Foundation (SNF; grants 31CA30_196644, 31CA30_196062, and 310030_173085), the European Commission (Marie Skłodowska-Curie Innovative Training Network “HONOURS”; grant agreement No. 721367, and the Horizon 2020 project “VEO” grant agreement No. 874735), the Federal Ministry of Education and Research (BMBF; grant RAPID, #01KI1723A) and by core funds of the University of Bern and the German Federal Ministry of Food and Agriculture. The hACE2-KI(B6) KI mice were generated with support from NIH/NIAID P01AI059576 (subproject 5) and partial support from NIH/NIAID U54-AI-057158. Partial support was from the US Centers for Disease Control and Prevention COVID-19 Task Force. We thank the next generation sequencing platform (NGS) of the University of Bern. We also acknowledge Mareen Lange, Christian Korthase, Michael Currier and Gloria Larson and Sandra Renzullo for their excellent technical assistance and Frank Klipp, Doreen Fiedler, Harald Manthei, Christian Lipinski, Jianzhong Tang, and Aurélie Godel for their invaluable support in the animal experiments.

This activity was reviewed by CDC and was conducted consistent with applicable federal law and CDC policy: 45 C.F.R. part 46, 21 C.F.R. part 56; 42 U.S.C. Sect. 241(d); 5 U.S.C. Sect. 552a; 44 U.S.C. Sect. 3501 et seq. The findings and conclusions in this manuscript are those of the authors and do not necessarily represent the official position of the US Centers for Disease Control and Prevention.

## Author contributions

VT, DW, MB, and CB conceived the study. TT, BZ, DH, AT performed most of the experiments. NE, SS, FL, JKe, HS, JP, HS, BT, JJ, RD, DH, NJH, LU, JKi, AP, BH, XF, XL, LW, NJ, DC, JH, MW, MK, TS, JR, SdB, CB conducted experimental work and/or provided essential experimental systems, data analysis, and reagents. VT, DW, MB, CB, TT, BZ, NE, SS, LT, DH, NJH, LU wrote the manuscript and made the figures. All authors read and approved the final manuscript.

## Data availability

Sequencing data from passage 1 virus stocks for recombinant SARS-CoV-2^S-614D^ and SARS-CoV-2^S-614G^, as well as data from all *in vitro* and *in vivo* competition experiments, will be made available on the NCBI Sequence Read Archive (SRA).

## Methods

### Cell and culture conditions

Vero E6 cells (ATCC) were cultured in Dulbecco’s modified Eagle’s medium (DMEM) supplemented with 10% fetal bovine serum, 1x non-essential amino acids, 100 units ml^−1^ penicillin and 100 μg ml^−1^ streptomycin. Baby Hamster Kidney cells expressing SARS-CoV N protein (BHK-SARS-N)^22^ were maintained in minimal essential medium (MEM) supplemented with 5% fetal bovine serum (FBS), 1x non-essential amino acids, 100 units ml^−1^ penicillin and 100 μg ml^−1^ streptomycin, 500 μg ml^−1^ G418 and 10 μg ml^−1^ puromycin. Twenty-four hours before electroporation, BHK-SARS-N cells were treated with 1 μg ml^−1^ doxycyclin to express SARS-CoV N protein. All cell lines were maintained at 37°C and in a 5% CO_2_ atmosphere.

### Recombinant proteins

Recombinant hACE2 protein (Cat: RP01266) was purchased from ABclonal. Recombinant SARS-CoV-2 proteins S1-614D and S1-614G with polyhistidine-tag (Cat: 40591-V08H, 40591-V08H3) were purchased from Sino Biological. All proteins were quantitated with Qubit Protein Assay (Thermo Fisher Scientific). SARS-CoV-2 S1-614D and S1-614G tagged with human IgG Fc fragment were constructed by insertion of the S1 region (residues 1-681) to pFUSE-hIgG1-Fc1 vector (InvivoGen, USA) and expressed using the Expi293 Expression system (Thermo Fisher Scientific). Supernatants were collected and quantified by western blotting using anti-human IgG secondary antibody (ThermoFisher A-21091). SARS-CoV-2 S-614D and S-614G proteins containing polyhistidine- and avi-tagged full-length ectodomain (residues 1-1208, furin cleavage site mutated) was also expressed using the Expi293 Expression system. The full-length ectodomain proteins were purified using HisTrap FF column (GE Life Sciences) in elution buffer containing 20 mM sodium phosphate, 0.5 M NaCl, 500 mM imidazole, pH 7.4, followed by desalting using Zeba spin desalting column (Thermo Fisher Scientific), per manufacturers’ instructions. The purified proteins were further concentrated on Amicon Ultra Centrifugal Filters (Sigma-Aldrich) and quantified using Qubit protein assay.

### Bio-Layer interferometry (BLI) assay

Affinity between human ACE2 to SARS-CoV-2 S1-614D, S1-614G, S-614D or S-614G were evaluated using Octet RED96 instrument at 30°C with a shaking speed at 1000 RPM (ForteBio). Streptavidin biosensors (SA) (ForteBio) were used. S1 proteins were pre-biotinylated using EZ-Link NHS-PEG4-Biotin (ThermoFisher Scientific). Following 20 minutes of pre-hydration of SA biosensors and 1 minute of sensor check, 100 nM S1-614D or S1-614G in 10X kinetic buffer (ForteBio) were loaded onto surface of SA biosensors for 5 minutes. After 2 minutes of baseline equilibration, 10 minutes of association was conducted at 2.5 to 75 nM of human ACE2, followed by 20 minutes of dissociation in the same buffer, which was used for baseline equilibration. S proteins with Avi-tag were pre-biotinylated using BirA biotin-protein ligase standard reaction kit (Avidity). 25 nM S-614D or 15 nM S-614G in 10X kinetic buffer (ForteBio) were loaded onto surface of SA biosensors for 5 minutes. After 2 minutes of baseline equilibration, 5 minutes of association was used for 2 to 32 nM hACE2, followed by 10 minutes of dissociation in 10X kinetic buffer. The data were corrected by subtracting reference sample, 1:1 binding model with global fit was used for determination of affinity constants.

### Flow cytometry

A stable clone of BHK cells expressing exogenous hACE2 were pelleted and resuspended in reaction buffer (PBS pH7.4 with 0.02% tween-20 and 4% BSA) at a concentration of 5 × 10^6^ cells/ml. 100 μl/well of the cells were aliquoted into a round-bottom 96-well plate and incubated on ice for at least 5 min. S1 proteins were diluted in reaction buffer on ice. 50 μl of S1 diluents were added into corresponding wells of cells and incubated on ice for 20 min with shaking. After incubation, cells were washed in 200 μl PBST washing solution (PBS pH7.4 with 0.02% tween-20) once and then 100 μl of 1:300 diluted secondary antibody (ThermoFisher Cat # A-21091 for Fc-tag and ThermoFisher Cat # MA1-21315-647 for polyhistidine-tag) was added into each well of cells, mixed, and incubated on ice with shaking for 15 min. After washing twice, cells were resuspended in 200 μl PBST and analyzed using the BD FACSCanto II Flow Cytometer. Data was processed with Flowjo_v10.6.1.

### Generation of infectious cDNA clones using TAR cloning and rescue of recombinant viruses

To introduce the 614G mutation to the Spike gene, PCR mutagenesis (Supplementary Table 1) was performed on the pUC57 plasmid containing SARS-CoV-2 fragment 10^10^ using Q5® Site-Directed Mutagenesis Kit (New England BioLab). Both D614 and G614 infectious cDNA clones were generated using in-yeast TAR cloning method as describe previously^10^. *In vitro* transcription was performed for EagI-cleaved YACs and PCR-amplified SARS-CoV-2 N gene using the T7 RiboMAX Large Scale RNA production system (Promega) as described previously^26^. Transcribed capped mRNA was electroporated into baby hamster kidney (BHK-21) cells expressing SARS-CoV N protein. Electroporated cells were co-cultured with susceptible Vero E6 cells to produce passage 0 (P.0) of the recombinant S-614D and S-614G viruses. Subsequently, progeny viruses were used to infect fresh Vero E6 cells to generate P.1 stocks for downstream experiments.

### Virus growth kinetics and plaque assay

Characterization of virus growth kinetics in Vero E6 was performed as described previously^10^. Twenty-four hours before infection, cells were seeded in a 24-well plate at a density of 2.0 × 10^5^ cells per ml. After washing once with PBS, cells were inoculated with viruses at multiplicity of infection (MOI) of 0.01. After 1 h, the inoculum was removed and cells were washed three times with PBS followed by supply with appropriate culture medium.

Plaque forming unit (PFU) per ml of recombinant S-614D and S614-G viruses were determined by plaque assay in a 24-well format. One day before infection, Vero E6 cells were seeded at a density of 2.0 × 10^5^ cells per ml. After washing once with PBS, cells were inoculated with viruses serially diluted in cell culture medium at 1:10 dilution. After 1 h of incubation, inoculum was removed, and cells were washed with PBS and subsequently overlaid with 1:1 mix of 2.4% Avicel and 2X DMEM supplemented with 20% fetal bovine serum, 200 units ml^−1^ penicillin and 200 μg ml^−1^ streptomycin. After 2 days of incubation at 37°C, cells were fixed in 4% (v/v) neutral-buffered formalin before stained with crystal violet.

Statistical significance was determined by two-sided unpaired Student’s *t*-test without adjustment for multiple comparisons.

### Infection of human nasal and bronchial epithelial cells

Primary human nasal epithelial cultures (hNE; MucilAir™ EP02, Epithelix Sàrl, Geneva, Switzerland) were purchased and handled according to the manufacturer instructions. Normal human bronchial epithelial (NhBE) cells were purchased (Emory University, Atlanta, GA, USA) and cultured according to recommended protocols. The hNE cultures were inoculated with 0.5×10^5^ PFU per well, or mixtures of 1:1, 10:1 and 1:10 of SARS-CoV-2^S-614D^ and SARS-CoV-2^S-614G^. NhBE cell cultures were inoculated with 1.0×10^5^ PFU per well, or with wild type isolates SARS-CoV-2/USA-WA1/2020 (USA-WA1, S-614D) or SARS-CoV-2/Massachusetts/VPT1/2020 (MA/VPT1, S-614G) at 2 x 10^5^ TCID_50_/well, For competition experiments, NhBE cells were inoculated with 1:1 or 9:1 mixed SARS-CoV-2^S-614D^ and SARS-CoV-2^S-614G^ at 1×10^5^ PFU per well. After incubation at 33°C for one or two hours, for hNE or NhBE cell cultures respectively, inoculum was removed, cells were washed, and subsequently incubated, as indicated, at 33°C, 37°C, or 39°C. To monitor viral replication, apical washes were collected every 24 hours. All titers were determined by standard plaque-assay or TCID_50_ on Vero E6 cells.

For competition experiments, viral RNA was extracted from apical washes using the QIAamp 96 Virus QIAcube HT Kit (QIAGEN). The SARS-CoV-2 genome was amplified using a highly multiplexed tiling PCR reaction based on the ARTIC protocol (https://www.protocols.io/view/ncov-2019-sequencing-protocol-bbmuik6w) with some modification. Briefly, primers were designed to produce overlapping 1kb amplicons (Supplementary Table 2). Reverse transcriptase was performed as described in the ARTIC protocol. The single cDNA reaction was carried forward by preparing two PCR reactions, one each for the odd and even pools of primers. Two primer pools (odds and evens) were prepared to contain 0.1 μM of each individual primer. Tiling PCR of the resultant cDNA was performed by combining 12.5 μL 2x Q5 polymerase, 5.5 μL water, 2 μL of the primer pool, and 5 μL of cDNA followed by incubating the reaction at 98°C for 30 seconds, 38 cycles of 98°C for 15 seconds and 63°C for 5 minutes, and holding at 4°C. Corresponding odd and even amplicons were normalized and pool for library preparation. Fragmented libraries were prepared using the Nextera XT DNA library preparation kit and sequenced via Illumina MiSeq. Analyses were performed using IRMA^27^ with a SARS-CoV-2 module.

### RNA extraction and RT-PCR

Preparation of viral RNA for next-generation sequencing was performed as described previously^10^. In brief, Vero E6 cells were infected with P.1 viruses. Extraction of total cellular RNA was done using Nucleospin^®^ RNA Plus kit (Macherey-Nagel) according to the manufacturers’ instruction.

RNA from hNE apical washes and mouse oropharyngeal swabs were prepared using QIAamp^®^ Viral RNA Mini Kit (QIAGEN) and Nucleospin^®^ RNA kit (Macherey-Nagel) according to the manufacturers’ protocol.

To determine the ratios of S-614D:S-614G in competition assays in Epithelix and hACE2-KI mice, reverse transcription PCR was performed on extracted RNA using SuperScript™ IV One-step RT-PCR System (Invitrogen). The sequence-specific primers were used to generate an amplicon of 905 bp covering the D614G region: 5’-AATCTATCAGGCCGGTAGCAC-3’ and 5’-CAACAGCTATTCCAGTTAAAGCAC-3’. RT-PCR reaction was performed in a 50-μl reaction using 0.01 pg to 1μg total RNA. The cycling parameters were set as follows: 50°C for 10 min, 98°C for 2 min; 35 cycles at 98°C for 10 sec, 55°C for 15 sec, and 72°C for 30 sec; and a 5-min incubation at 72°C. DNA concentration was determined using Qubit dsDNA HS (High Sensitivity) Assay (Thermo Fisher), and subsequently diluted to 200 ng in 50 μl of nuclease-free water for sequencing by Nanopore sequencing MinION.

RNA extraction and preparation of RT-PCR reactions were performed in low- and no-copy laboratory environment to avoid contamination.

### Sequencing and computational analysis

Recombinant SARS-CoV-2^S-614D^ or SARS-CoV-2^S-614G^ RNAs of P.1 stock were sequenced by next-generation sequencing as described previously^10^. Briefly, RNA was extracted from Vero E6 cells infected with either recombinant SARS-CoV-2^S-614D^ or SARS-CoV-2^S-614G^ using the NucleoSpin RNA Plus kit (Macherey-Nagel) according to the manufacturer’s guidelines. Sequencing libraries were subsequently prepared using the Illumina TruSeq Stranded mRNA Library Prep kit (Illumina, 20020595) and pooled cDNA libraries were sequenced across two lanes on a NovaSeq 6000 S1 flow cell (2 x 50bp, 100 cycles) using the Illumina NovaSeq 6000 platform (Next Generation Sequencing Platform, University of Bern, Switzerland). For data analysis, TrimGalore v0.6.5 was used to remove low-quality reads and adaptors from the raw sequencing files. The resulting trimmed paired-end reads were then aligned to the SARS-CoV-2 genome (GenBank accession MT108784) using Bowtie2 v2.3.5. Finally, a consensus sequence was generated for each virus stock using Samtools v1.10 with the -d option set to 10,000. Data analysis was performed on UBELIX, the HPC cluster at the University of Bern (http://www.id.unibe.ch/hpc).

### Animal studies

#### Ethics declarations

The hACE-2 knock-in mice were originally generated at the Wadsworth Center, New York State Department of Health IACUC protocol # 09-405 (Wentworth, PI). Mouse experimentation was conducted in compliance with the Swiss Animal Welfare legislation and animal studies were reviewed and approved by the commission for animal experiments of the canton of Bern, Switzerland under license BE-43/20. All of the ferret and hamster experiments were evaluated by the responsible ethics committee of the State Office of Agriculture, Food Safety, and Fishery in Mecklenburg-Western Pomerania, Germany (LALLF M-V), and gained governmental approval under registration number LVL MV TSD/7221.3-1-041/20. This project also obtained clearance from the CDC’s Animal Care and Use Program Office.

#### Human ACE2 knock-in mouse study

Generation of hACE2 knock-in mice. The hACE2-KI(B6) (B6.Cg-*Ace2^tm1^*^(ACE2)Dwnt^/J) line was generated by targeted mutagenesis to express human ACE-2 cDNA in place of the mouse *Ace2* gene. Thus, in this new animal model, hACE2 expression is regulated by the endogenous mouse *Ace2* promoter/enhancer elements. The targeting vector (WEN1-HR) had hACE2 cDNA inserted in frame with the endogenous initiation codon of the mouse *Ace2* (Extended Data Figure 2A). The human cDNA was flanked by an FRT-neomycin-FRT-loxP cassette and a distal loxP site. WEN1-HR was used to transfect 129Sv/Pas ES cells and 837 ES cell clones were isolated and screened for homologous recombination by PCR and Southern blot. Eleven properly recombined ES cell clones were identified and some of them were used for blastocyst injection and implantation into female mice to generate 22 male founders with chimerism (129Sv/Pas:C57BL/6) ranging from 50 to 100%. To complete the hACE2 knock-in, we crossed the chimeric males with C57BL/6J Flp-expressing females to excise the FRT flanked neomycin selection cassette and generate the floxed humanised ACE2 allele (Extended Data Figure 2A). These hACE2 knock-in mice were identified by PCR and confirmed by Southern Blot and were backcrossed to C57BL/6J mice for 7 generations (N7) prior to this study. This line has been donated to The Jackson Laboratory for use by the scientific community (Stock 035000).

Heterozygous hACE2-KI female mice were obtained from The Jackson Laboratory (USA) and C57BL/6J wild-type (WT) female mice were from Janvier Lab (France). All mice were acclimatized for at least 2 weeks in individually ventilated cages (blue line, Tecniplast), with 12/12 light/dark cycle, autoclaved acidified water, autoclaved cages including food, bedding and environmental enrichment at the SPF facility of the Institute of Virology and Immunology, Mittelhäusern, Switzerland. One week before infection, mice were placed in individually HEPA-filtered isolators (IsoCage N, Tecniplast). Mice (10-12-week-old) were anesthetized with isoflurane and infected intranasally with 20μl (i.e., 10μl per nostril) with a 1:1 ratio of the SARS-CoV-2 variants (WT and hACE2-KI mice) or mock culture medium (WT mice only). After intranasal infection, mice were monitored daily for body weight loss and clinical signs. Throat swabs were collected daily under brief isoflurane anesthesia using ultrafine sterile flock swabs (Hydraflock, Puritan, 25-3318-H). The tips of the swabs were place in 0.5 mL of RA1 lysis buffer (Macherey-Nagel, Ref. 740961) supplemented with 1% β-mercaptoethanol and vortexed. Groups of mice from two independent experiments were euthanized on days 2 and 4 p.i. and organs were aseptically dissected avoiding cross-contamination. Systematic tissue sampling was performed: (1) lung right superior lobe, right nasal concha, right olfactory bulb, part of the right brain hemisphere, apical part of the right kidney, parts of the distal small intestine (ileum) were collected for RNA isolation in RA1 lysis buffer; (2) lung middle, inferior and post-caval lobes, left nasal concha, left olfactory bulb, part of the right brain hemisphere, part of the right kidney were collected in MEM; (3) lung left lobe, liver left lobe, left kidney, left brain hemisphere and part of the jejunum and ileum were fixed in buffered formalin. Data were generated from two identically designed independent experiments.

Mouse tissue samples collected in RA1 lysis buffer supplemented with 1% β-mercaptoethanol were homogenized using a Bullet Blender Tissue Homogenizer (Next-Advance). Homogenates were centrifuged for 3 min at 18,000 g and stored at −70°C until processing. Total RNA was extracted from homogenates using the NucleoMag VET kit for viral and bacterial RNA/DNA from veterinary samples (Macherey Nagel, Ref: 744200) and the KingFisher Flex automated extraction instrument (ThermoFisher Scientific) following manufacturers’ instructions. RNA purity was analyzed with a NanoDrop 2000 (ThermoFisher Scientific). A 25 μl RT-PCR for the viral E gene was carried out using 5 μl of extracted RNA template using the AgPath-ID One-Step RT-PCR (Applied Biosystems). Samples were processed in duplicate. Amplification and detection were performed in a Applied Biosystem 7500 Real-Time PCR Systems under the following conditions: an initial reverse transcription at 45 °C for 10 min, followed by PCR activation at 95 °C for 10 min and 45 cycles of amplification (15 seconds at 95 °C, 30 seconds at 56 °C and 30 seconds at 72 °C). Relative quantification of virus load in swabs and mouse tissues was based on β2 microglobulin expression measured by TaqMan qPCR according to the manufacturer’s protocol (Assay mM00437762_m; ThermoFisher).

Fixed mouse tissue samples were processed, sectioned and stained with hematoxylin and eosin (H&E) at the COMPATH core facility (University of Bern). Histopathological lung slides of hACE2-KI mice and wild-type mice (infected and mock) were examined and scored by a board-certified veterinary pathologist (SdB), who was blinded to the identity of the samples. Scoring criteria are detailed in Extended Data Table 2.

#### Hamster study

Six Syrian hamsters, *Mesocricretus auratus*, (Janvier Labs, France) were infected intranasally under a short-term inhalation anesthesia with 70 μl of SARS-CoV-2^S-614D^ and SARS-CoV-2^S-614G^ at equal ratios using 10^4.77^ TCID_50_/animal (calculated from back titration of the original material). After 24 hours of isolated housing in individually ventilated cages (IVCs), six pairs, each with one directly inoculated donor hamster and one sham-inoculated contact hamster were co-housed. The housing of each hamster duo was strictly separated in individual cage systems to prevent spill-over between different pairs. For the following seven days (day 2 until day 8 after infection) and on day 12 after infection, viral shedding was monitored in addition to a daily physical examination and body weighting routine.

To evaluate viral shedding, nasal washes were individually collected from each hamster under a short-term isoflurane anesthesia. Starting with the pair’s contact hamster, each nostril was rinsed with 100 μl PBS (1.0x phosphate-buffered saline) and reflux was immediately gathered. A new pipet tip for every nostril and hamster was used to prevent indirect spill-over transmission from one animal to another. Furthermore, in-between the respective pairs, a new anesthesia system was used for each pair of animals. At day 8 post infection, one contact hamster was found dead. Although having lost up to almost 20% of their body weight, every other hamster recovered from disease. Under euthanasia, serum samples were collected from each hamster.

For a second experimental setup seven hamsters each were infected via the intranasal route with 10^5.1^ TCID_50_/animal of SARS-CoV-2^S-614D^, or 10^4.5^ TCID_50_/animal SARS-CoV-2^S-614G^ (calculated from back titration of the original material). From day 1 onwards to day 4 nasal washes were obtained from these hamsters and body weight changes recorded. A tissue panel from respiratory organs, including nasal conchae, tracheal tissue, cranial, medial, and caudal portions of the right lung, and the pulmonary lymph node, were collected after euthanasia from these hamsters.

#### Ferret study

In accordance with the hamster study, twelve ferrets (*Mustela putorius furo*) from in-house breeding, were divided into six groups of two individuals. Ferret pairs were of equal age between four and 18 months. In total, four ferrets were female (two pairs) and eight ferrets were male or neutered male (four pairs). The housing of each ferret duo was strictly separated in individual cage systems, to prevent spill-over between different pairs. Per separate cage, one individual ferret was inoculated intranasally, by instillation of 125 μl of the aforementioned inoculum (1×10^5^ TCID_50_/animal; calculated from back titration of the original material) into each nostril under a short-term isoflurane-based inhalation anesthesia. Time points and sampling technique corresponded to the methods used for the hamsters, albeit ferret nasal washing was performed using two times 700μl PBS per animal. The contact-ferret was not inoculated.

#### Specimen work-up, viral RNA detection, sequencing and quantification analyses

Organ samples were homogenized in a 1 ml mixture of equal volumes composed of Hank’s balanced salts minimum essential medium (MEM) and Earle’s balanced salts MEM containing 2 mM l-glutamine, 850 mg l-1 NaHCO3, 120 mg l-1 sodium pyruvate and 10% FCS (supplemented with 10% Fetal Calf Serum and 1% penicillin–streptomycin) at 300 Hz for two minutes using the Tissuelyser II (Qiagen, Hilden, Germany) and centrifuged to clarify the supernatant. Nucleic acid was extracted from 100 μl of the nasal washes after a short centrifugation step or organ sample supernatant using the NucleoMag Vet kit (Macherey Nagel, Düren, Germany). Viral loads in these samples were determined using the nCoV_IP4 RT-PCR^28^ including standard RNAs that where absolute quantified by digital droplet PCR with a sensitivity of 10 copies/reaction. The extracted RNA was afterwards subjected to MinION sequencing and digital droplet PCR. For MinION sequencing, amplicons were produced with primers (WU-21-F: AATCTATCAGGCCGGTAGCAC; WU-86-R: CAACAGCTATTCCAGTTAAAGCAC) using SuperScript IV One-step RT-PCR (Thermo Fisher Scientific; Waltham, MA USA). Amplicons were sequenced on a MinION system using from Oxford Nanopore using Native Barcoding 1-12 (EXP-NBD104) and 13-24 (EXP-NBD114), respectively with Ligation Kit SQK-LSK109 (Oxford Nanopore; Oxford, UK). Libraries were loaded on R9.4.1 Flow Cells (FLO-MIN106D) on a MinION coupled to a MinIT. Realtime high accuracy basecalling, demultiplexing and barcode and adapter trimming was performed with MinKnow v20.06.17, running Guppy vs4.0.11. Downstream analysis was done in Geneious 2019 vs2.3. Read length filtered eliminated reads < 800 and > 1100 nt and remaining reads were mapped in subsets of 10,000 reads to the amplicon reference undergoing two iterations, with custom sensitivity allowing a maximum of 5% gaps and maximum mismatch 30%. Variants were analyzed on specific position with calculating p-values for every variant. Ratio fraction A/G was calculated from numbers of reads as fraction=Areads/(Areads+Greads).

For rare mutation and sequence analysis based on digital PCR, the QX200 Droplet Digital PCR System from Bio-Rad (Hercules, CA, USA) was used. For RT-PCR, the One-Step RT-ddPCR Advanced Kit for Probes (Bio-Rad) was applied according to the supplier’s instructions. For the amplification of an 86 bp fragment of both variants of the spike protein gene, the forward primer SARS2-S-1804-F (5’-ACA AAT ACT TCT AAC CAG GTT GC-3’) and the reverse primer SARS2-S-1889-R (5’-GTA AGT TGA TCT GCA TGA ATA GC-3’) were used. For the detection of the D and G variants in one sample, two allele specific locked nucleic acid (LNA) probes were applied: SARS2-S-v1D-1834FAM (5’-FAM-TaT cAG gat GTt AAC-BHQ1-3’) and SARS2-S-v3G-1834HEX (5’-HEX-T cAG ggt GTt AAC-BHQ1-3’). The LNA positions are depicted in lower case. The concentration of the primers and probes was 20μM and 5μM, respectively. For data analysis, the QuantaSoft Analysis Pro software (version 1.0.596) was used.

**Extended Data Fig. 1. Additional *in vitro* characterization of S proteins and SARS-CoV-2^S-614D^ and SARS-CoV-2^S-614G^ isolates.**

**(a)** Affinity between S and hACE2 determined by Bio-layer interferometry. Biotinylated spike protein (ectodomain) (S-614D or S-614G) was loaded onto surface of streptavidin biosensors and association was conducted using hACE2 followed by dissociation. Data represent two biological replicates. (**b)** Binding of Fc-tagged or polyhistidine-tagged S1 to BHK-hACE2 cells determined by flow cytometry. Mean Fluorescence intensity is shown for corresponding S1 protein concentration. Data represent the mean ± s.d of three biological replicates. (**c)** Replication of wild type SARS-CoV-2/USA-WA1/2020 (S-614D) and SARS-CoV-2/Massachusetts/VPT1/2020 (S-614G) isolates in NhBE cells at 33°C (left), 37°C (middle), and 39°C (right). NhBE cells were infected with 200,000 TCID50 of each virus. Supernatants were collected every 24 h and titrated by TCID50 assay. Data represent the mean ± s.d. of four replicates. Statistical significance was determined by two-sided unpaired Student’s *t*-test without adjustments for multiple comparisons. *P* values (from left to right): left, NS *P* = 0.7874; *\*P* = 0.0328; NS *P* = 0.1887; NS *P* = 0.8985; NS *P* = 0.5296; middle, NS *P* = 0.1475; NS *P* = 0.1415; ** *P* = 0.0033; NS *P* = 0.3184; right, NS *P* = 0.6018; NS *P* = 0.3903; NS *P* =0.0898; \**P* = 0.0445. (**d)** Competition assay of recombinant SARS-CoV-2^S-614D^ and SARS-CoV-2^S-614G^ in NhBE at 33°C, 37°C and 39°C. The inoculum was prepared by mixing two viruses at 9:1 ratio based on plaque forming unit ml^−1^. NhBE was infected with 100,000 pfu of the 9:1 virus mix. Viral RNA was extracted from daily supernatant and sequenced by next generation sequencing. Bar graph shows proportion of sequencing reads encoding either S-614D or S-614G, and each square dot represents one individual data point.

**Extended Data Figure 2. hACE-2 KI mouse generation and infection with SARS-CoV-2 ^−614D^ and SARS-CoV-2^S-614G^ viruses.** (**a**) Design and generation of humanized ACE2 knock-in mice (hACE2-KI). The hACE2 cDNA was inserted in frame with the endogeneous initiation codon of mouse Ace2 in exon 2, which was deleted. The hACE2 cDNA was flanked 5’ with a loxP site (black triangle) and 3’ with a FRT-neomycin-FRT-loxP cassette. The targeting construct included a negative selection cassette (PGK-DTA) to improve selection of clones with homologous recombination. Chimeric male mice transmitting the targeted locus were crossed with Flp-deleter female mice to generate the floxed hACE2 knock-in allele. This allele can be used : (1) without further cre-mediated recombination, as shown here, to study humanized ACE2 mice (hACE2-KI), where the hACE2 cDNA is expressed in place of mouse Ace2; (2) after crossing with a cre-deleter mouse line to generate constitutive *Ace2* knock-out mice; (3) after crossing with tissue-specific cre line. Ubiquitous and tissue-specific knock-out mice can be crossed with conventional hACE2 transgenic mice to remove the endogenous mouse ACE2, which could confound pathogenesis studies that may occur due to heterodimerization of ACE2. (**b**) Body weight loss at indicated time points after infection of hACE2-KI mice (n=8), wild-type mice infected (n=9) and for mock-infected wild-type sampled identically (n=10). (**c**) Quantitative

RT-PCR analysis of tissue homogenates of inoculated hACE2-KI and wild-type mice at indicated time points.

**Extended Data Figure 3. Virus replication in infected Syrian hamsters.**

**(a)** Body weight loss at indicated time points after infection of Syrian hamsters. Donor animals (n=6; black dots) were intranasally inoculated with SARS-CoV-2^S-614D^ / SARS-CoV-2^S-614G^ at equal ratio (1×10^4.77^ TCID_50_/animal as determined by backtitration of the original inoculum). 24h after infection, naïve hamsters (n=6; orange triangles) were housed in direct contact in an “one-to-one” experimental setup. **(b)** Quantitative RT-PCR analysis of individual nasal washing samples obtained from donor hamsters and contact animals, respectively. **(c)** Body weight loss at the indicated time points after infection of the hamsters. Syrian hamsters were inoculated with 10^5.1^ TCID_50_/animal of SARS-CoV-2^S-614D^ (n=7, blue dots), or 10^4.5^ TCID_50_/animal SARS-CoV-2^S-614G^ (n=7, red triangles) via the intranasal route. Titers were determined by backtitration of the original inoculation material. **(d)** Viral genome copy numbers are shown as determined by RT-qPCR from individual nasal washing samples of the animals inoculated with the single variant virus. **(e)** Quantitative RT-PCR analysis of tissue homogenates of inoculated hamsters of the SARS-CoV-2^S-614D^ group (n=7, blue dots) versus the SARS-CoV-2^S-614G^ group (n=7, red triangles).

**Extended Data Figure 4. “Twin”-inoculation of donor ferrets with equal ratios of SARS-CoV-2^S-614D^ and SARS-CoV-2^S-614G^.**

Donor ferrets (black dot; n=6) were intranasally inoculated with 10^5.4^ TCID_50_/animal as determined by back titration of an inoculum comprising equal ratios of SARS-CoV-2^S-614D^ and SARS-CoV-2^S-614G^. Twenty four hours post inoculation one contact ferret (orange triangle; n=6) was commingled with one donor ferret, creating six donor – contact ferret pairs. **(a)** Individual body weight of ferrets at the indicated days, relative to the day of inoculation, is plotted. **(b)** Genome copy numbers for inoculated donor and contact ferrets. Individual nasal washing samples of the indicated days were analyzed by RT-qPCR “nCoV_IP4”, and absolute numbers were calculated using a set of standard RNAs. All donor ferrets (black dots) tested vRNA positive, starting already day 2 post inoculation (n=6). 4 out of 6 contact ferrets (orange triangles) tested vRNA positive beginning with day 4 (corresponding with day 3 after contact). 2 of the 6 contact ferrets never tested positive for vRNA throughout the study.

**Extended Data Table 1 | Lung histopathological score of mice infected with SARS-CoV-2.** Data is shown for individual hACE2-KI mice (K1-K8) and wild-type inoculated (WT1-WT9) and wild-type mice mock inoculated (M1-M10). Scoring criteria are detailed in Extended Data Table 2. Scoring was performed by a pathologist blinded sample identification.

**Extended Data Table 2 | Score sheet of lung histopathology.**

## References

1 Korber, B. et al. Tracking Changes in SARS-CoV-2 Spike: Evidence that D614G Increases Infectivity of the COVID-19 Virus. Cell 182, 812–827.e819, doi:10.1016/j.cell.2020.06.043 (2020).

2 Zhu, N. et al. A Novel Coronavirus from Patients with Pneumonia in China, 2019. N Engl J Med 382, 727–733, doi:10.1056/NEJMoa2001017 (2020).

3 Zhou, P. et al. A pneumonia outbreak associated with a new coronavirus of probable bat origin. Nature 579, 270–273, doi:10.1038/s41586-020-2012-7 (2020).

4 ECDC. COVID-19 pandemic, <https://www.ecdc.europa.eu/en/covid-19-pandemic> (2020).

5 Huang, C. et al. Clinical features of patients infected with 2019 novel coronavirus in Wuhan, China. Lancet 395, 497–506, doi:10.1016/s0140-6736(20)30183-5 (2020).

6 Letko, M., Marzi, A. & Munster, V. Functional assessment of cell entry and receptor usage for SARS-CoV-2 and other lineage B betacoronaviruses. Nature Microbiology 5, 562–569, doi:10.1038/s41564-020-0688-y (2020).

7 Wrapp, D. et al. Cryo-EM structure of the 2019-nCoV spike in the prefusion conformation. Science 367, 1260–1263, doi:10.1126/science.abb2507 (2020).

8 Hoffmann, M. et al. SARS-CoV-2 Cell Entry Depends on ACE2 and TMPRSS2 and Is Blocked by a Clinically Proven Protease Inhibitor. Cell 181, 271–280.e278, doi:10.1016/j.cell.2020.02.052 (2020).

9 Yurkovetskiy, L. et al. Structural and Functional Analysis of the D614G SARS-CoV-2 Spike Protein Variant. Cell, doi:10.1016/j.cell.2020.09.032 (2020).

10 Thi Nhu Thao, T. et al. Rapid reconstruction of SARS-CoV-2 using a synthetic genomics platform. Nature 582, 561–565, doi:10.1038/s41586-020-2294-9 (2020).

11 Wu, F. et al. A new coronavirus associated with human respiratory disease in China. Nature 579, 265–269, doi:10.1038/s41586-020-2008-3 (2020).

12 Bao, L. et al. The pathogenicity of SARS-CoV-2 in hACE2 transgenic mice. Nature 583, 830–833, doi:10.1038/s41586-020-2312-y [pii]10.1038/s41586-020-2312-y [doi] (2020).

13 Jiang, R. D. et al. Pathogenesis of SARS-CoV-2 in Transgenic Mice Expressing Human Angiotensin-Converting Enzyme 2. Cell 182, 50–58 e58, doi:S0092-8674(20)30622-X [pii]10.1016/j.cell.2020.05.027 [doi] (2020).

14 Sia, S. F. et al. Pathogenesis and transmission of SARS-CoV-2 in golden hamsters. Nature 583, 834–838, doi:10.1038/s41586-020-2342-5 (2020).

15 Imai, M. et al. Syrian hamsters as a small animal model for SARS-CoV-2 infection and countermeasure development. 117, 16587–16595, doi:10.1073/pnas.2009799117 %J Proceedings of the National Academy of Sciences (2020).

16 Osterrieder, N. et al. Age-Dependent Progression of SARS-CoV-2 Infection in Syrian Hamsters. Viruses 12, doi:v12070779 [pii] viruses-12-00779 [pii] 10.3390/v12070779 [doi] (2020).

17 Imai, M. et al. Syrian hamsters as a small animal model for SARS-CoV-2 infection and countermeasure development. Proc Natl Acad Sci U S A 117, 16587–16595, doi:10.1073/pnas.2009799117 (2020).

18 Richard, M. et al. SARS-CoV-2 is transmitted via contact and via the air between ferrets. Nat Commun 11, 3496, doi:10.1038/s41467-020-17367-2 (2020).

19 Kim, Y. I. et al. Infection and Rapid Transmission of SARS-CoV-2 in Ferrets. Cell Host Microbe 27, 704–709.e702, doi:10.1016/j.chom.2020.03.023 (2020).

20 Alouane, T. et al. Genomic Diversity and Hotspot Mutations in 30,983 SARS-CoV-2 Genomes: Moving Toward a Universal Vaccine for the “Confined Virus”? Pathogens 9 (2020).

21 Zhang, J. et al. Structural impact on SARS-CoV-2 spike protein by D614G substitution. 2020.2010.2013.337980, doi:10.1101/2020.10.13.337980 %J bioRxiv (2020).

22 Shi, P. Y. et al. Spike mutation D614G alters SARS-CoV-2 fitness and neutralization susceptibility. Res Sq, doi:rs.3.rs-70482 [pii]10.21203/rs.3.rs-70482/v1 [doi] (2020).

23 Zhang, L. et al. The D614G mutation in the SARS-CoV-2 spike protein reduces S1 shedding and increases infectivity. bioRxiv, doi:2020.06.12.148726 [pii]10.1101/2020.06.12.148726 [doi] (2020).

24 Li, Q. et al. The Impact of Mutations in SARS-CoV-2 Spike on Viral Infectivity and Antigenicity. Cell 182, 1284–1294 e1289, doi:S0092-8674(20)30877-1 [pii]10.1016/j.cell.2020.07.012 [doi] (2020).

25 van den Worm, S. H. et al. Reverse genetics of SARS-related coronavirus using vaccinia virus-based recombination. PLoS One 7, e32857, doi:PONE-D-11-21011 [pii]10.1371/journal.pone.0032857 [doi] (2012).

26 Thiel, V., Herold, J., Schelle, B. & Siddell, S. G. Infectious RNA transcribed in vitro from a cDNA copy of the human coronavirus genome cloned in vaccinia virus. J Gen Virol 82, 1273–1281, doi:10.1099/0022-1317-82-6-1273 [doi]> (2001).

27 Shepard, S. S. et al. Viral deep sequencing needs an adaptive approach: IRMA, the iterative refinement meta-assembler. BMC Genomics 17, 708, doi:10.1186/s12864-016-3030-6 (2016).

28 Pasteur, I. Protocol: Real-time RT-PCR assays for the detection of SARS-CoV-2. WHO Document (accessed 10/21/2020): https://www.who.int/docs/default-source/coronaviruse/real-time-rt-pcr-assays-for-the-detection-of-sars-cov-2-institut-pasteur-paris.pdf?sfvrsn=3662fcb6_2 (2020).

